# Analysis of antibiotic response in Clinical Wound *Pseudomonas aeruginosa* isolates: Unveiling Proteome Dynamics of tobramycin tolerant phenotype

**DOI:** 10.1101/2024.04.16.589491

**Authors:** Kasandra Buchholtz, Rosa Jersie-Christensen, Karen Angeliki Krogfelt, Biljana Mojsoska

## Abstract

*Pseudomonas aeruginosa* (*P. aeruginosa*) is an opportunistic human pathogen, causing serious chronic infections. *P. aeruginosa* can adapt efficiently to antibiotic stressors via different genotypic or phenotypic strategies such as resistance and tolerance. The adaptation regulatory system is not always very well understood. In this study, we use shotgun proteomics to investigate the system-level response to tobramycin in two clinical wound *P. aeruginosa* isolates and PAO1. We profiled each strain for its antibiotic drug-tolerant phenotype using supra-minimum inhibitory concentrations (supra-MIC) of tobramycin and applied proteomics to investigate the protein expression profiles. The MIC revealed that all isolates were susceptible to tobramycin but at supra-MIC concentrations at stationary growth, a degree of tolerance was observed for the isolates. We identified around 40 % of the total proteins encoded by the *P. aeruginosa* genome and highlighted shared and unique protein signatures for all isolates. Comparative proteome profiling in the absence of antibiotic treatment showed divergent fingerprints, despite similarities in the growth behavior of the isolates. In the presence of tobramycin, the isolates shared a common response in the downregulation of proteins involved in the two-component system, whereas stress response proteins were present at higher levels. Our findings provide insight into the use of proteomic tools to dissect the system-level response in clinical isolates in the absence and presence of antibiotic stress.

## Introduction

Wound infections caused by bacteria are a major health concern globally, especially in immunocompromised patients, but not only such as those suffering from diabetes ^1,2^. Infection of the wound usually starts with bacterial colonization, and when colonization is combined with factors such as decreased vascular supply and host immune defense, it can become an actual infection. The microbiology of chronic wounds is genuinely complex, and it is often necessary to take quantitative biopsies of the infected tissue to determine the course of treatment ^3^. The opportunistic pathogen *Pseudomonas aeruginosa* (*P. aeruginosa*) is the most frequently recovered bacteria in clinical specimens ^4^. It is a major nosocomial pathogen with increasing relevance to human health and disease, particularly in chronic wound infections in diabetic and hospitalized patients. *P. aeruginosa* infections are extremely challenging to treat because of the high intrinsic and acquired antibiotic resistance ^5^ and also due to their ability to alternate between planktonic and biofilm lifestyles. However, recent studies have shown the presence of a minor, transiently multidrug-tolerant subpopulation of cells, with a clinical emphasis on their role in recurrent infections ^6^. *P. aeruginosa* tolerant population is a dormant variant of the antibiotic-susceptible population that becomes transiently tolerant to antimicrobial agents in the presence of the agent ^7,8^. These types of cells tolerate high concentrations of antibiotics, even as high as 100x the reported minimum inhibitory concentrations ^8,9^. The size of the drug-tolerant subpopulation within a bacteria population is believed to be controlled by stress signaling pathways, such as the stringent response (SR), SOS, and stress response ^7,10^. With the development of novel assays and new omics technologies (proteomics, metabolomics, transcriptomics, etc.) to investigate the effect of antibiotics, we now know that some, such as tobramycin ^11^ and polymyxin ^12^ can have dual effects that are concentration-dependent, and these effects can further be different depending on the bacterial growth phase. At high concentrations both agents can cause substantial damage to the outer membrane and tobramycin also inhibits protein synthesis by blocking translation ^13,14^. Full proteome studies have been used to investigate the system-level response of dynamic changes in bacterial species in the presence of antibiotics ^15^. However, very few attempts have been made to use proteomics to characterize the drug-tolerant phenotype ^16^, characterize core proteome in differently grown bacteria ^17^ and investigate the effects of antibiotics such as tobramycin on planktonic *P. aeruginosa* ^18^ as well as *P. aeruginosa* biofilms ^19^. In this study, we use tobramycin, an antibiotic that belongs to the group of aminoglycoside antibiotics, with dual activity. And look at system-level proteome response in clinical isolates and PAO1 at stationary growth. Previous transcriptome studies have shown large differences in gene expression in the PAO1 genome when comparing actively grown bacteria and those in stationary growth phase ^20^, however, no studies have reported on proteome response at stationary grown bacteria in MHBII media. Our study provides useful insights in this area, that can be further used to expand our understanding of the utilization of proteomics to investigate and revisit antibiotics’ mode of action on a system level.

### Experimental Procedures

#### Bacterial strains and growth conditions

The bacterial strains used were clinical wound isolates of *P. aeruginosa* that had been collected from two patients admitted to Copenhagen Wound Healing Centre, Bispebjerg Hospital in Denmark with venous leg ulcers in 2001, (Patient 1: 9000a) and (Patient 2: 989a). The wound isolates were collected using filter paper pads and charcoal swabs throughout eight weeks of regular wound observations. The clinical wound isolates were then processed at the State Serum Institute (SSI). *P. aeruginosa*, PAO1 WT was used for comparison. MHBII media and LB agar plate were used to culture *P. aeruginosa*.

#### Antibiotic tolerance profiling

At the stationary phase, the culture of PAO1, 9000a, and 989a was subjected to (4 µg/ml) tobramycin. The concentration was chosen to be at least 10x the minimum inhibitory concentration (MIC). The survival of clinical wound isolates and PAO1 was determined based on the assessment of cell viability after 24 hours of antibiotic exposure. A serial dilution of the cultures was performed in sterile saline and 10 µl were plated on LB agar plates. Colony forming units per ml (CFU/ml) were counted. Experiments were performed with three biological replicates (n=3). Data is shown in the supplementary material.

#### Killing curve and antibiotic tolerance assay validation

For validation of survival, a time-kill experiment was performed in parallel to proteomic sample preparation. Bacteria cultures were grown in MHBII media. Antibiotic-treated (4 µg/ml tobramycin) samples were harvested at several time points (0h, 6h, 24h, and 48h), together with the corresponding control of untreated bacteria. The samples were centrifugated at 14000 rpm for 10 minutes, washed with ice-cold 0.9% w/v NaCl (ACS reagent ≥ 99%, Honeywell), and a serial dilution of the cultures was performed in saline after which 10 µl were plated on LB agar plates and incubated overnight at 37°C, followed by survival assessment (CFU/ml, Figure S1).

#### Proteomic Sample preparation

To generate a drug-tolerant subpopulation for the proteomic analysis, the overnight culture of 9000a, 989a, and PAO1 was diluted 1:100 in 40 ml pre-heated MHBII and incubated overnight at 37°C, 180 rpm, to reach the stationary phase. The sample for time zero was harvested, centrifugated at 14000 rpm for 10 minutes, and washed with cold 0.9% w/v NaCl (ACS reagent ≥ 99%, Honeywell). The bacteria cultures were then split into quartets, with a corresponding control. The culture was subjected to 4 µg/ml tobramycin and incubated for 24 hours at 37°C,180 rpm. After 24 hours of exposure, a sample was harvested, centrifuged at 14000 rpm for 10 minutes, was washed with cold 0.9% w/v NaCl. For reproducibility, four biological and four technical replicas were used in this experiment. The cell pellet was resuspended and washed 2x with ice-cold PBS. The pellet was resuspended in 200 µl pre-heated lysis buffer (6M Guanidine hydrochloride (GuHCl), 100mM Tris-HCl pH 8.5, 5mM tris(2-carboxyethyl)-phosphine (TCEP) and 10mM Chloroacetamide (CAA)) and incubated for 10 minutes at 99°C. After incubation, the lysate was sonicated for 1 minute. The protein concentration was then measured on Nanodrop. The samples were diluted with 25 mM Tris-HCL pH 8.5 to a final protein concentration of 200 µg/ml. Trypsin (Thermo Scientific, Pierce) was added in a 1:50 w/w enzyme: protein ratio, and the samples were digested overnight at 37°C at 550 rpm. After digestion the reaction was stopped by adding 10% Trifluoroacetic acid (TFA) to a final concentration of 1% and the samples were centrifuged at 14000 rpm for 5 minutes. Sep-Pak C18 columns (Water Corporation, Milford, MA) were washed with 1x with 100% acetonitrile (ACN) and 2x with 0.1% (TFA). After centrifugation, the supernatant was loaded onto the Sep-Pac C18 column, washed 2x with 0.1% (TFA), and eluted in two steps, using 40% and 60% (ACN). SpeedVac concentrator was used to concentrate the peptide samples. Approximately 1µg was loaded onto the LC-MS/MS instrument.

#### LC-MS/MS

Peptides were trapped on a C18 column (5 μm, 5mm, 0.3mm) and separated on a 15 cm fused silica column (75 μm inner diameter) pulled and packed in-house with 1.9 μm C_18_ beads (Reprosil-AQ Pur, Dr.

Maisch) on an Ultimate 3000 system connected to a LTQ Velos Orbitrap (Thermo Scientific, San Jose, US). The peptides were separated with a 110 min gradient with increasing buffer B (90% ACN and 0.1% formic acid), going from 5 to 30% in 70 min, 30 to 50% in 15 min, 50 to 95% in 20 min followed by a 5 min wash and re-equilibrating step. All steps were performed at a flow rate of 250 nL/min. The LTQ Velos Orbitrap was operated in data-dependent top 15 mode. Full scan mass spectra were recorded in the orbitrap at a resolution of 60,000 at *m*/*z* 200 over the *m*/*z* range 375–1600 with a target value of 1 × 10^6^ and a maximum injection time of 500 ms. CID-generated product ions were recorded in the ion trap with a maximum ion injection time set to 100 ms and a target value set to 1 × 10^4^. The spray voltage was set to 2.2 kV, S-lens RF level at 50, and heated capillary at 300 °C. Normalized collision energy was set at 35 and the isolation window was 2 *m*/*z*.

#### Proteomic data analysis

All raw LC-MS/MS data files were processed using MaxQuant version 1.6.5.0 (Cox & Mann, 2008) with default settings and LFQ for quantification. The data was searched against entries matching organism ID 208964 in the UniProt database, https://www.uniprot.org/, downloaded in June 2019. Using 1% FDR at both peptide and protein levels. Perseus software (version 1.6.14.0), was used for further processing of the data. A principal components analysis (PCA) was performed to determine the quality of the samples, together with a multiple comparison test, to visualize the differences in protein expression. For the protein quantitative analysis, a student t-test was conducted, and differentially expressed proteins were filtered based on the following cut-off; false discovery rate (FDR) of 0.05 and log2-Fold Change of 2 (s0=2). Hierarchical cluster analysis was performed based on the significance found in the student t-test. The functional enrichment annotation and visualization were performed using online tools DAVID (version 2021) and DeepVenn^21^. Protein identifications were filtered at a false discovery rate (FDR) of 0.05. To further highlight the potentially important proteins (enrichment terms) among the differentially expressed proteins, STRING (version 12.0) was used to predict and visualize protein-protein interactions. KEGG pathway and pseudomonas genome database ^22^, were used for further functional annotation of proteins. Information about all up and down-regulated proteins in the tobramycin-treated samples is provided in Supplementary files (Excel files 1-9). The mass spectrometry proteomics data have been deposited to the ProteomeXchange Consortium via the PRIDE^23^ partner repository with the dataset identifier PXD051528.

#### Statistical analysis

In the validation of the antibiotic-tolerant assay, the viability was tested to estimate the survival % after the bacterial culture had been exposed to tobramycin for 24 hours. Significance was assessed using a two-tailed student t-test with unequal variances in log-transformed values and a p-value threshold of 0.05. FDR and s0:2 whenever applied. The statistical analysis was performed and visualized in Excel/ Prism (GraphPad 10.0.0).

## Results and discussion

### Tobramycin tolerant bacterial population was observed in all 3 isolates at supra-MIC concentrations

In our study, we used high concentrations of tobramycin (supra-MIC, 10 x MIC) to investigate the effect on bacteria in stationary phase growth of three *P. aeruginosa* strains in MHBII media, and the survival profiles were assayed (Figure 1A). The samples at 24 hrs were collected for treatment for proteomic analysis (Figure 1B). One (989a) out of the 3 strains was shown to have a high tobramycin-tolerant phenotype (Figure S1) by measuring viability on plates.

**Figure 1.**
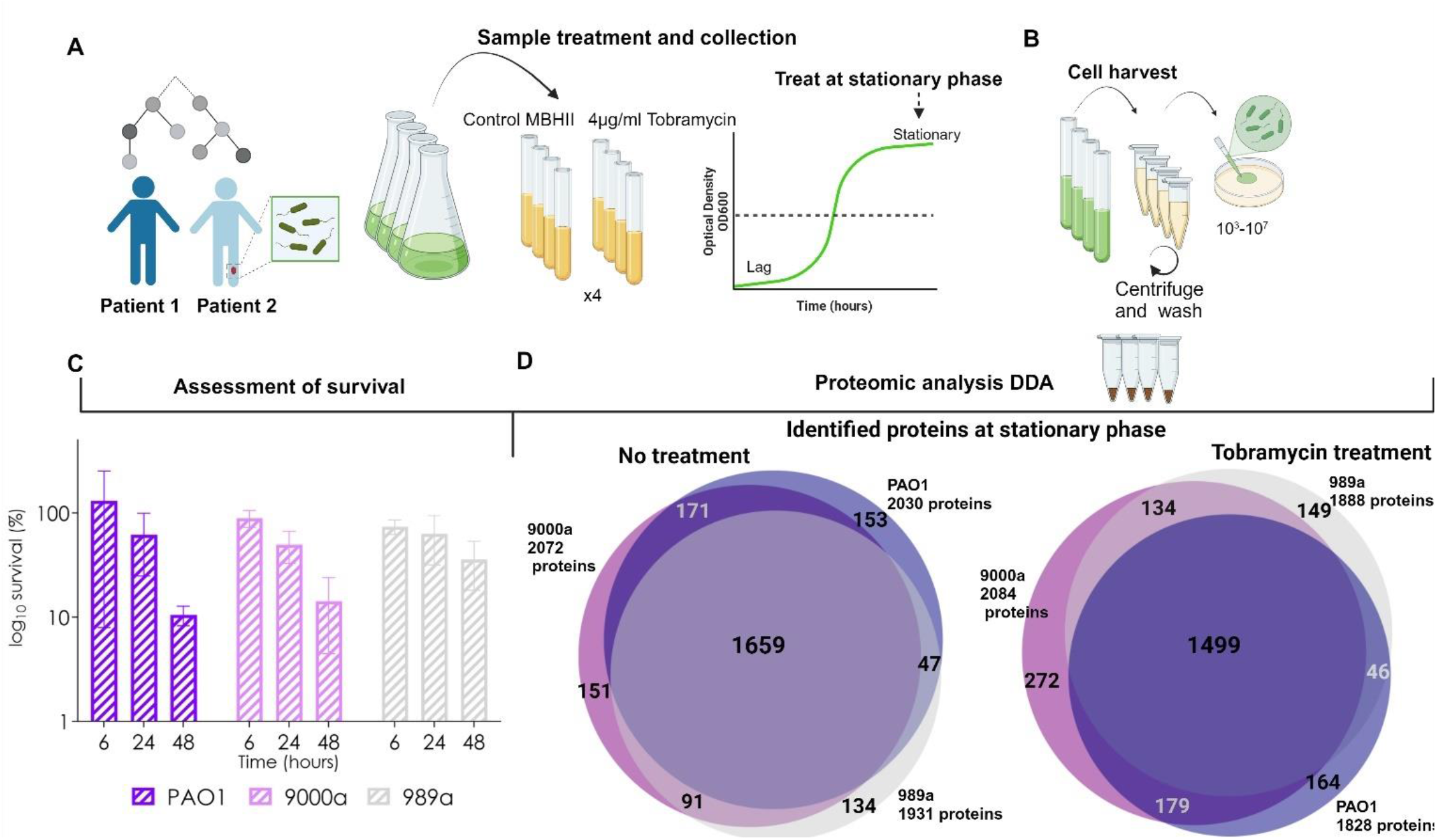
Schematic representation of the experiment workflow. A) The killing assay was performed on PAO1 and two clinical wound isolates 989a, 9000a. The overnight culture was diluted 1:100 in 40 ml pre-heated MHBII and incubated for 24 hours at 37 C°, 180 rpm. The culture was split into two groups, untreated, and treated samples with 4 µg/ml tobramycin, followed by incubation further for 24 hours at 37C°, 180 rpm. B) 1 ml of each culture was harvested at time points 0h (control samples) and after 24h of treatment (treated samples). The culture was centrifuged, washed in the cold 0.9% w/v NaCl, and plated on the LB agar plate to determine CFU/ml for validation of sub-population of drug tolerant cells. The pellet was saved at -20ºC for proteomic sample preparation. C) Time dependent bacterial killing of stationary growth PA isolates treated with supra-MIC concentrations of Tobramycin (4 ug/ml). Data normalized to the control of untreated bacteria. two-way ANOVA was conducted to compare tobramycin impact on the cell survival between isolates and the time point. No significant difference between the isolates PAO1, 9000a (patient 1), and 989a (patient 2) (F(2)=0.16, P<0.85) was observed but a significant difference in the cell survival over time (F(3)=4.630, P<0.01) was present. No significant difference in the overall interactions (F(6)=0.3, P<0.93) is observed. D) Venn diagrams^21^ showing the total number of identified proteins for all three PA strains and the number of shared and unique proteins for each isolate at stationary growth phase after treatment with 4 ug/ml tobramycin (left). Protein Ids were used for comparison. Created with Biorender.com

**Figure 1**.

A time-kill assay of both clinical isolates and PAO1 at 6, 24, and 48 hrs shows different survival profiles after antibiotic exposure (Figure 1C). Tobramycin is a bactericidal antibiotic that inactivates the initiation complex of translation ^13,14^. Tobramycin most often acts on metabolically active cells (cells in early to mid-exponential growth), to impair active protein synthesis through ribosome binding. However, our data shows that tobramycin effectively kills bacteria in stationary growth when compared with the untreated population, and we associate this activity with the drug tolerance profile ^24^ of the tested *P. aeruginosa* strains. The antibiotic effect in all strains is most significant over time, where there is a continuous growth inhibition observed (Figure 1C). Tobramycin killing curve patterns is different among the 3 investigated PA strains (9000a, 989a, and PAO1) with a pronounced time effect on bacterial survival for PAO1 and 9000a (p<0.01) and less for 989a (Figure 1C).

### Proteome coverage in PAO1 and two clinical wounds PA isolates at the stationary phase

Proteomic analysis of system response level on the mode of action of tobramycin in MHB II media has not been studied before. We therefore collected bacterial pallets of stationary grown cells that were untreated and treated with supra-MIC concentrations of tobramycin for 24 hours of growth. The samples were analyzed using DDA, shotgun proteomics approach. To date, there are 5,570 annotated Open Reading Frames in PAO1 (6.3 million base pairs, PseudoCAP^25^). We could identify more than 2000 proteins (∼40 % of all proteins) in the *P. aeruginosa* isolates (PAO1, 989a, and 9000a) used in the present study in MH II media (Figure 1D). LFQ intensities were used as protein abundances, and these were correlated for a specific strain and time point. We used PCA (not shown) to visualize variance among the replicates and experimental conditions. We found that the biological replicates cluster with one another and antibiotic-treated ones were separated from the control (data not shown).

### Unique and shared protein responses in P. aeruginosa isolates at stationary growth phase

Previous studies on the characterization of transcriptomes in PAO1 in stationary phase planktonic culture in LB media revealed that 19.4 % of the PAO1 genome is differently expressed between log phase and stationary phase and the profile of confluent biofilms were related to those of a stationary phase grown bacteria. Here genes that are involved in general stress (quorum-sensing or RpoS genes), glycogen accumulation and breakdown, and oxidative stress (*katE, katN*) have been characterized ^26^. When we compared the proteome profiles of all isolates between each other, at stationary growth, we found that many of the proteins *(n=1659*, Figure 1D) were shared, pointing out similarities in adaptation at this growth phase. As a control, we could identify markers of the stationary growth phase such as RpoS, a sigma factor molecule that is a central regulator in stationary phase ^27^, to be expressed in all isolates (data not shown). Functional annotation clustering analysis (David, version 2021) showed proteins that are associated with translation, to have the highest enrichment score (data not shown). We could identify a few unique proteins for each isolate (Figure 3). These unique proteins were analyzed for functional annotation clustering (biological processes and molecular function) using David annotation tool ^28^ (not shown). Many of the proteins were representative of the lipopolysaccharide biosynthetic process as well as metabolic processes covering carbohydrate synthesis and lipids for PAO1 (n=153), small molecule synthesis, and RNA metabolic process for 9000a (n=151) and enzyme and receptor activity, as well as processes involved in cellular responses and regulation of gene expression among others (n=134). This is the first study to report the full proteome profiling in MHBII media at stationary phase planktonic growth for these isolates without any antibiotic treatment, and as such it allows us to use this information for further comparative studies.

### Unique and shared protein responses in P. aeruginosa isolates with Tobramycin treatment

All isolates were treated with tobramycin at stationary phase for 24 hrs and the proteome profiles were again compared, leading to identification of 1499 shared proteins between all three isolates. From those identified, we could observe a greater similarity for 9000a and 989a (n=179, Figure 1D). Several unique proteins were also identified and their profiles and top unique proteins with enrichment analysis are presented in Figures S2-4).

### Single-strain proteome analysis

#### Tobramycin growth inhibition of stationary PA cells -effect of supra-MIC concentrations

Previous data have shown that at the mid-exponential growth phase, a tobramycin concentration lower than 4 µg/ml inhibits protein synthesis, and a concentration higher than 8 µg/ml kills the bacteria through membrane disruption ^29^. These expression patterns could indicate that tobramycin acts as a protein synthesis inhibitor, which triggers a misfolded oxidized protein that can form potential toxin aggregates, lethal to the bacterial cell, rather than killing through membrane disruption ^30^. To look at individual strain response of supra-MIC concentrations of tobramycin on a proteome level, we compared protein expressions of treated and untreated samples of each isolate at stationary phase at 24 hrs. Our single strain comparative proteomic analysis for PAO1, 9000a, and 989a, revealed several hundreds of differentially expressed proteins in response to tobramycin treatment. In PAO1, 285 proteins were differentially expressed from which 112 proteins were up- and 173 proteins downregulated (Figure 2 A). The heat map shows great clustering based on the biological origin of the samples. Functional enrichment analysis of the up- and down-regulated proteins revealed that, in response to tobramycin stress, proteins belong to 3 enrichment classes covering multiple biological and molecular processes such as transport (cell membrane proteins, antibiotic resistance, stress response (cytoplasmic proteins, protein folding, etc.) (Figure 2B). Five protein clusters were enriched in the down-regulated group of proteins, and these belong to metabolic pathways, amino acid metabolism, metabolic processes (biosynthesis of amino acids), and proteins in the two-component system (Figure 2C). For each analysis, differentially expressed proteins (DEPs) (FC>1.5, p<0.05) were cross-checked for shared identification or isolate-specific response. Unique proteins are presented with arrows. Protein FpvA is uniquely expressed as an adaptive response in PAO1 with appx. 4.2 x increased expression in the tobramycin-treated sample. This protein is a ferripyoverdine receptor and has a versatile role in the transport of small molecules (Pseudomonas Genome Data Base). Some of the top downregulated proteins in the antibiotic-treated samples highlight processes involved in the two-component system (unknown, unique target proteins (PA2573, PA2920, and PA4520)), metabolic pathways (fatty acid, amide biosynthesis, unique proteins PA0744, LiuC, LiuB, AmbE, PvdP, PvdF, PchF, and ArcA), cellular metabolic processes (unique proteins MetK, PhzB2, PA2330, and GlyA2) as well as protein biosynthesis (Figure 5 B-D).

**Figure 2.**
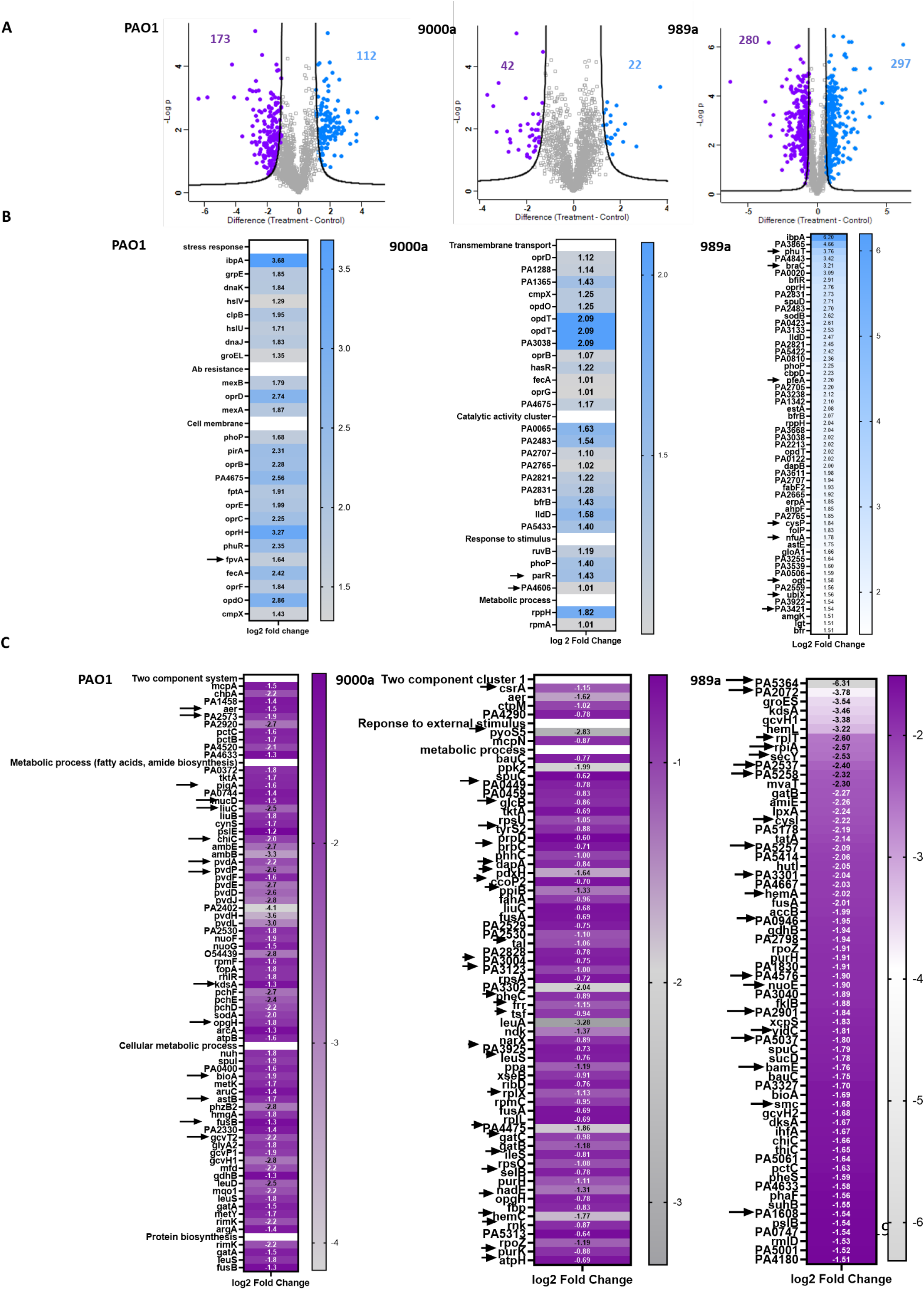
A) Volcano plot showing differentially expressed proteins (DEPs) proteins in isolates PAO1, 9000a, and 989a in the antibiotic-treated samples. 285 proteins were significantly changed in PAO1 threshold FDR 0.05, s:0=2, (n=3). Volcano plot showing 64 differentially expressed proteins (DEPs) in the antibiotic-treated isolates 9000a, FDR 0.05, s:0=2, (n=4). Volcano plot showing 577 differentially expressed proteins with statistical significance in the antibiotic-treated isolate 989a, FDR 0.05, s:0=2, (n=4). B) Heat map of log 2-Fold Change of selected upregulated proteins (FC>1.5) in all 3 isolates. Arrows show unique proteins. Proteins belong to stress response, antibiotic resistance, and cell membranes. C) Heat map of log 2-Fold Change of selected downregulated proteins (FC>1.5) in all 3 isolates. Arrows show unique proteins.

In 9000a there were 64 proteins differently expressed proteins (Figure 2A) between the tobramycin-treated and untreated samples. The proteins associated with higher expression patterns were clustered as transmembrane proteins and proteins that belong to metabolic processes (Figure 2B, unique proteins shown with arrows). Enrichment analysis of the downregulated proteins showed many to be associated with the two-component system, response to external stimulus, and various metabolic processes (Figure 2C, unique proteins shown with arrows). Proteins encoded by genes *parR* (two-component system response regulator) and PA4606 (conserved hypothetical proteins) were uniquely found to be upregulated in this strain when compared to the others. Two-component signal transduction systems enable bacteria to sense, respond, and adapt to changes in their environment or their intracellular state. Each two-component system consists of a sensor protein-histidine kinase and a response regulator. Two-component pathways thus often enable cells to sense and respond to stimuli by inducing changes in transcription ^31^. In the list of downregulated proteins multiple proteins were unique and protein PyoS5 (defense response) had the highest down expression level of 7 times less than in the control. This is an interesting finding, as pyocins act as bacteriocins towards other gram-negative species and if tobramycin downregulated production of this in vitro, then it causes decreased virulence and increased susceptibility for this strain.

**Figure 2**.

In isolate 989a, many proteins (n= 577), were found to be differentially expressed (Figure 2A, FDR=0.05, log2 FC, s0=2.). Proteins involved in the transport of small molecules, transcriptional regulators, porin activity, and biofilm were among those found enriched (Figure 2B, unique proteins shown with arrows). Among these proteins, PhuT (ABC transporter), BraC (amino acid transport), and PfeA (two-component system) had the highest translational level when compared to the control. A total of 64 proteins had significantly lower expression in the tobramycin-treated samples and among these 20 were unique for the 989a strain (Figure 2C, unique proteins shown with arrows).

### Significant DEPs shared among all isolates

The general understanding of the bacteria’s cellular response to tobramycin is that to survive, the bacteria will try to counteract the effect of the antibiotic, reducing the membrane permeability and enhancing ribosomal changes ^32^. A recent study has reported that the effect of tobramycin on the translation is reduced through enhanced gene expression, involved in ribosomal rescue, tRNA modification, and type II toxin-antitoxin (TA) system in *P. aeruginosa* ^33^. Interestingly, it was discovered through functional enrichment analysis that some of the upregulated proteins in this study were translation initiation factors (InfC, InfB), ribosomal proteins (RplO, RplX, RpsA, RpsO), and a transferase (RimK). Despite similar findings, it is not easy to compare the study of Sesso et al. because of the experimental differences. They exposed the cells at mid-exponential to 64 µg/ml, a relatively high tobramycin concentration, and used RNA and Ribosomal sequencing ^33^. In our study with 4 ug/ml tobramycin treatment of stationary planktonic cells, we have found that translational initiation factors were not among the DEPs in all 3 strains ((InfB, PAO1 (log2FC=-0.1), 9000a (log2FC=-0.4), 989a (log2FC=-0.45), (InfC, PAO1 (log2FC=0.48), 9000a (log2FC=-0.48), 989a (log2FC=0.12)). Concerning the ribosomal proteins, only RpsO, RplX, and RpsA were identified with their respective fold changes ((RpsO, PAO1 (log2FC=0.1), 9000a (log2FC=-1), 989a (log2FC=-1.2), (RplX, PAO1 (log2FC=0.035), 9000a (log2FC=-1.1), 989a (log2FC=-1.3)),(RpsA, PAO1 (log2FC=-0.39), 9000a (log2FC=-0.71), 989a (log2FC=-0.79)), (RimK, PAO1 (**log2FC=-2.23**), 9000a (log2FC=-0.2), 989a (log2FC=-0.74)) (Supplementary excel files 1-3). Only transferase RimK was found to be significantly downregulated in PAO1 upon treatment with high concentrations of tobramycin.

**Figure 3**.

**Figure 3.**
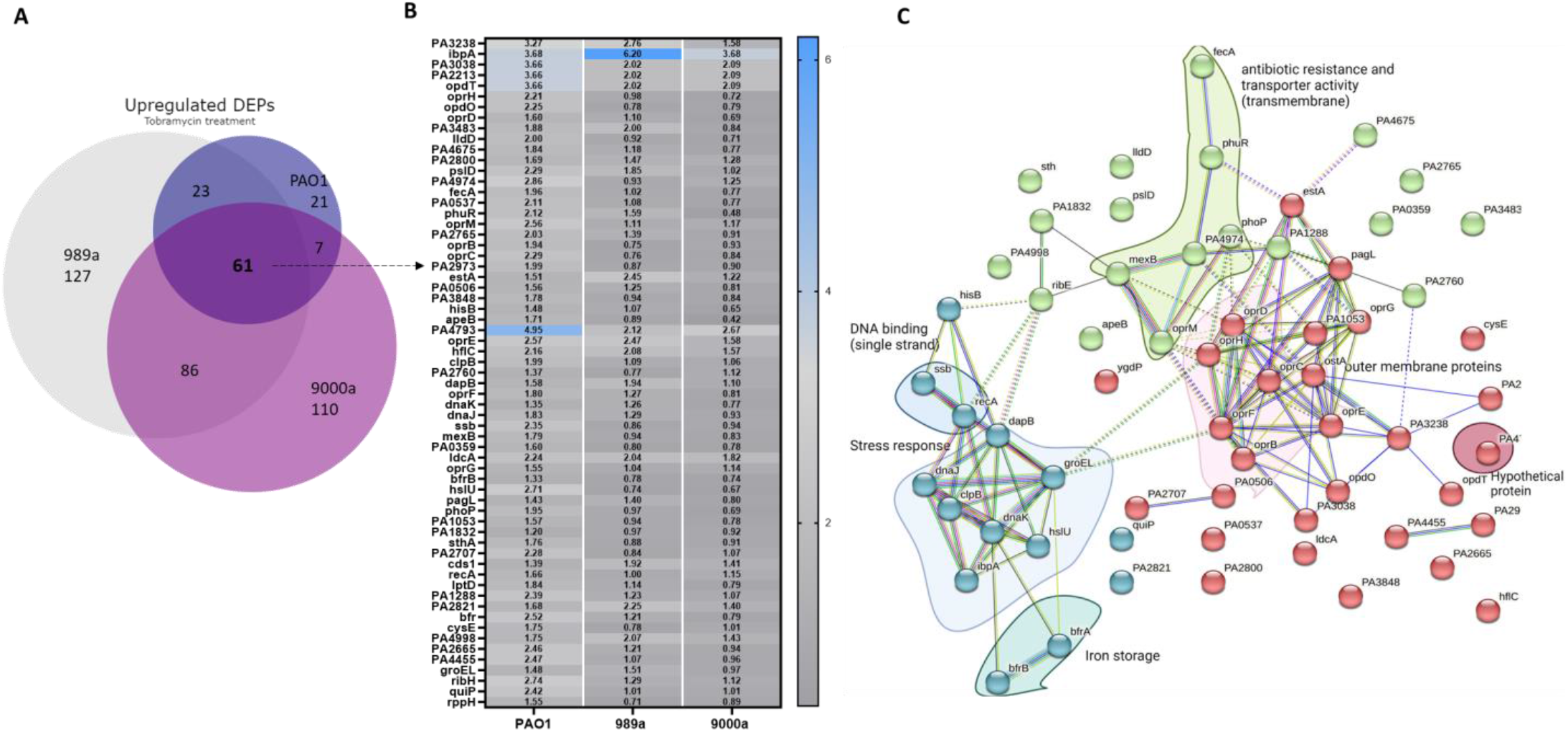
A) Venn diagram of all differentially expressed proteins (DEPs) that are upregulated in the antibiotic-treated P. aeruginosa isolates, PAO1, 989a, and 9000a. B) Log2 Fold Change of 61 commonly shared upregulated DEPs. C) Visualization and enrichment of the 61 proteins was done using STRING ( version 12.0)

From all upregulated proteins in the tobramycin-treated *P. aeruginosa* strains, we found that 61 proteins were found to be shared between all isolates, 86 proteins between the wound isolates, and 23 proteins between PAO1 and 989a. The log 2-Fold Change of the shared upregulated proteins is shown in Figure 3. Even though all the 3 isolates showed shared responses to tobramycin, the level of protein expression of the same proteins was different, supporting the different levels of survival pattern seen in Figure 1C. In PAO1, the hypothetical protein PA4793 exerts the highest expression levels under tobramycin stress conditions. In another case, the heat-shock protein IbpA involved in protein folding is seen to be expressed at a higher degree in all strains, particularly in 989a (Figure 3). When compared with the proteome changes in published proteome studies, we found that this protein is 90-fold more expressed in tobramycin-treated *P. aeruginosa* by Wu 2015. Subsequent deletion of this protein did not however increase the sensitivity of the strain towards tobramycin, but double mutants ibpA/ X, ibpA/ Y, and ibpA/HslV did increase the sensitivity to the antibiotic ^34^. In our study, we did find other heat shock proteins, previously reported, such as hslU among the topmost expressed proteins in all strains as well as the molecular chaperone protein DnaK ^34^. Concerning the most downregulated proteins enriched in all strains, the membrane protein IptF had the lowest expression levels by more than 60-fold when compared with the control (Figure 4). STRING enrichment analysis for the functional relationship of proteins that are downregulated in the tobramycin-treated samples also showed proteins that belong to the two-component system as well as proteins found in metabolic processes and translational elongation to be important for the adaptation of all PA strains to the stress response to tobramycin.

**Figure 4:**
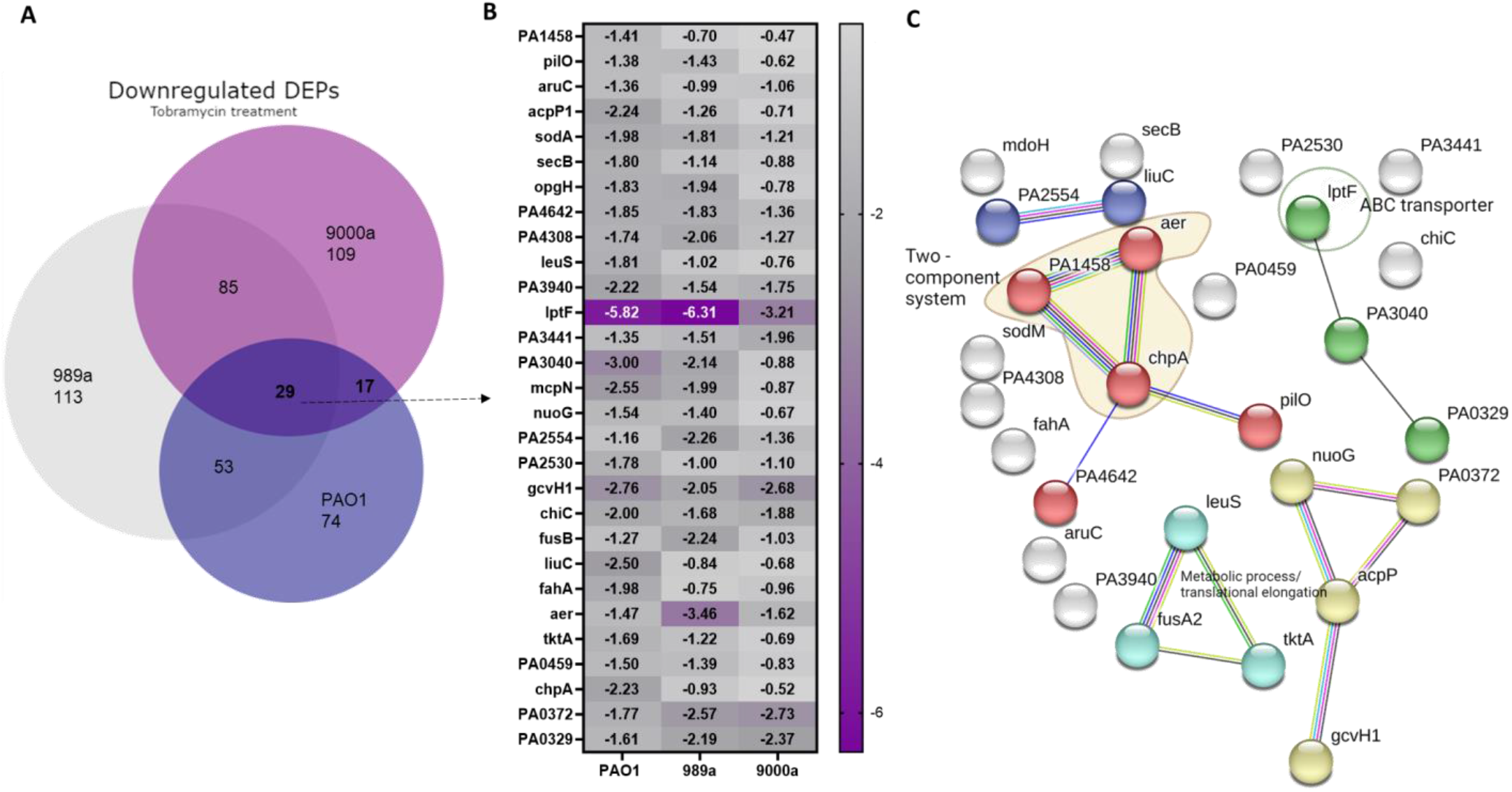
A) Venn diagram of all differentially expressed proteins (DEPs) that are down-regulated in the antibiotic treated P. aeruginosa isolates, PAO1, 989a and 9000a. B) Log2 FC of the 29 commonly shared downregulated DEPs C) Visualization and enrichment of the 29 proteins was done using STRING ( version?). Most proteins are membrane associated proteins.

**Figure 4**.

In the effort to characterize more proteins that could be specific among the strains, we approached two by two strain comparisons, and a few metabolic and hypothetical proteins LeuA, PvdH (PAO1 vs 9000a), PpiD, PA5061 (989a vs 9000a), and PA3690, PchG, PchE, PA2789 (PAO1 vs 989a) showed to exert expression profile specific for the specific strain pairing (Figure 5).

**Figure 5**.

**Figure 5:**
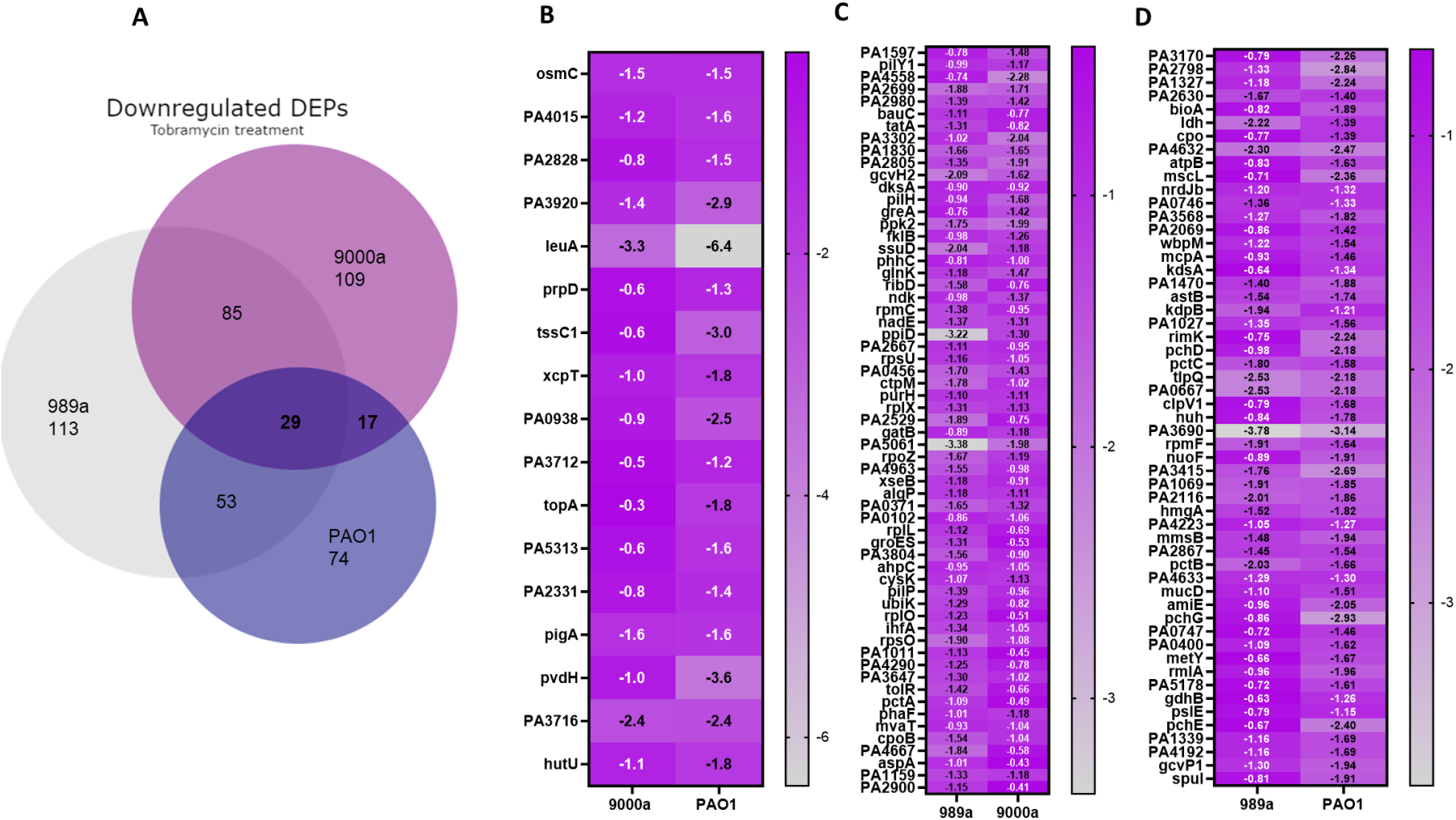
A) Venn diagram of all shared differentially expressed proteins (DEPs) that are downregulated in the antibiotic-treated P. aeruginosa isolates, PAO1, 989a, and 9000a. Log2 FC of shared DEPs between isolate-pairs, B) 9000a-PAO1, C) 989a-9000a, and D) 989a-PAO1.

Furthermore, a study presented by Viducic et al.^35^ reported that RpoS, the alternative sigma factor, is induced during the stationary phase, has a role in antibiotic tolerances, and postulated that the increased level of RpoS could be a consequence of the accumulation of ppGpp ^35^. We identified RpoS in all isolates, however, it did not appear to be significantly changed for the tobramycin-treated samples (Supplementary material). Additionally, published studies have discovered that DksA mutant could survive exposure to quinolones, postulating that DksA might act as a negative regulator on gene expression in putative genes involved in antibiotic tolerance ^35^. We found this protein to be downregulated in PAO1 (log2FC=-1.17), 9000a (log2FC=-0.929 and 989a (log2FC=-0.9). In addition to our comparative studies, we investigated the fold change of the most common proteins involved in virulence, antibiotic resistance, and biofilm, and their respective fold change are shown in Figure 5. Reduced susceptibility to aminoglycosides has been previously demonstrated in *P. aeruginosa* fusA mutant library ^36^. In our studies, protein expression of this fusA1 is downregulated in all strains by 0.5-fold when compared to the untreated control (Figure 6). All the strains had tobramycin MIC of 0.4 ug/ml which renders them susceptible to the drug. Therefore, small downregulation of this protein does not seem to play a role in the observed drug tolerance phenotype at stationary phase with 10-fold higher concentrations of the aminoglycoside. With regards to the efflux pump system MexAB-OprM, also reported to be relevant for aminoglycoside resistance when in high expression ^36^, our data supported the findings as MexAB-OprM proteins were all upregulated to a different extent between the isolates, with the highest degree in PAO1 (log2FC =1.8-2.3) (Figure 6). This finding did not correlate with the observed viability of PAO1 for wound isolates, as at 24 hrs they all exhibit similar degrees of survival. However, in 989a, there was a higher upregulation of the transcriptional regulator MexT where the protein was downregulated in PAO1. MexT mutants have previously been reported to show reduced expression of the membrane protein OprD ^37^, however, here we report significantly increased expression of protein OprD, despite the MexT levels (Figure 6). It is important to note that genotype-phenotype studies are not always a good predictor for the correlation of protein-protein interaction and multiple genotypic and phenotypic changes are needed for adaptation to the environment ^37^. Together with our study, the overall literature findings support the hypothesis that there is a dynamic and divergent proteome response in *P. aeruginosa* which is concentration and growth phase-dependent.

**Figure 6.**
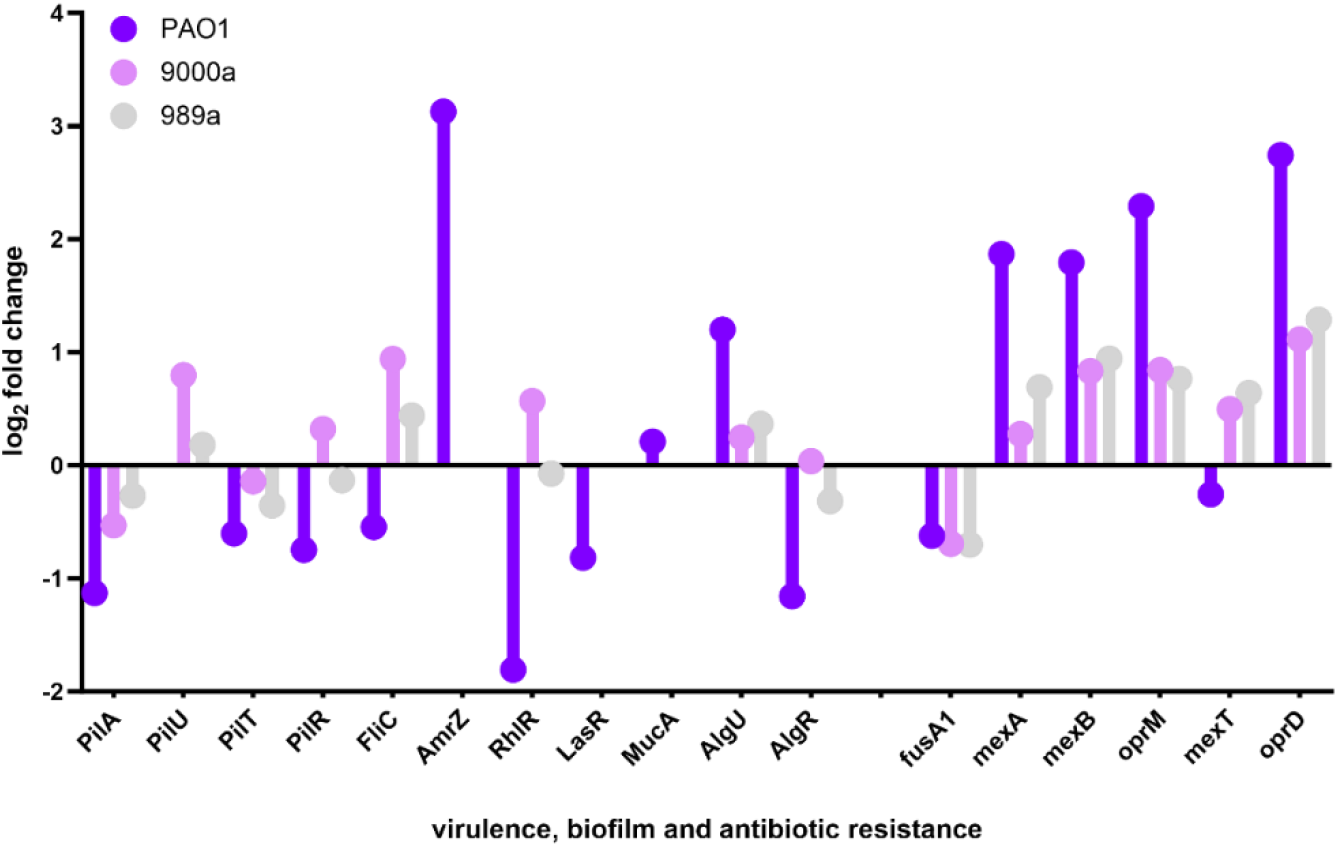
Proteome response in 3 clinical isolates, PAO1, 9000a, and 989a of proteins involved in virulence, biofilm formation, and antibiotic resistance. Log 2-Fold Change of all proteins is shown.

**Figure 6**.

## Conclusion and perspectives

This study aimed to investigate the proteome profile of 2 clinical wound isolates of *P. aeruginosa* and PAO1 in the presence of supra-MIC concentrations of tobramycin, through label-free shotgun proteomics and explore their antibiotic-tolerant phenotype. This is the first study that reports proteome profiling in MHBII media and therefore the findings are difficult to compare with others as a selection of growth media is an important consideration of proteomic experiments and can lead to approximately 10 % alternation ^38^ in the overall reported findings. We show that tobramycin exerts a time-dependent killing and affects the protein expression profiles of all 3 isolates at the stationary phase. Our findings point out that to survive, bacteria counteract the antibiotic effect by enhancing gene expression of proteins involved in ribosomal rescue, which supports the general understanding of the bacteria’ cellular response to antibiotic exposure. Comparative proteomics analysis with previously published data is difficult as not every study had the same experimental conditions as our data. On the protein level, PAO1, and the two wound isolates shared some significantly expressed proteins, hence showing a similarity in adaptation. In addition, the three-sequence types also had unique proteins, indicating that the response to tobramycin was not identical. The proteome profiling demonstrates that proteomics can be used to explore the differences in growth response between bacterial strains as well as characterize system-level responses to tobramycin, where actual proteins can be identified. The overall findings lead to generating a comprehensive understanding of the biological system interplay in investigating antibiotic responses and different phenotypes such as the tolerant phenotype. It is still challenging to interpret proteomic data and understand the impact of the system analysis, however, this study should motivate further comparative analysis and support computational efforts to investigate antibiotic responses in *P. aeruginosa*. To enhance our understanding of the observed response, future studies could investigate the time as a descriptor in addition to using the pulse labeling technique to identify potential protein targets.

## Supporting information

Table S1 Figure S1-4 Excel files 1-9

## Author information Corresponding Author Biljana Mojsoska

Associate Professor in Chemical Biology, Department of Science and Environment, Roskilde University, Denmark, Universitetsvej 1, 4000, Roskilde, *email: biljana@ruc*.*dk*

## Notes

The authors declare no competing financial interest.

## Author Contributions

Writing original draft BM, KJP

Final writing and editing BM

Data analysis BM, RJC, KJP

Data generation: KJP

All authors approve the final version of the document.

---

**Table of Content (TOC) Figure:**
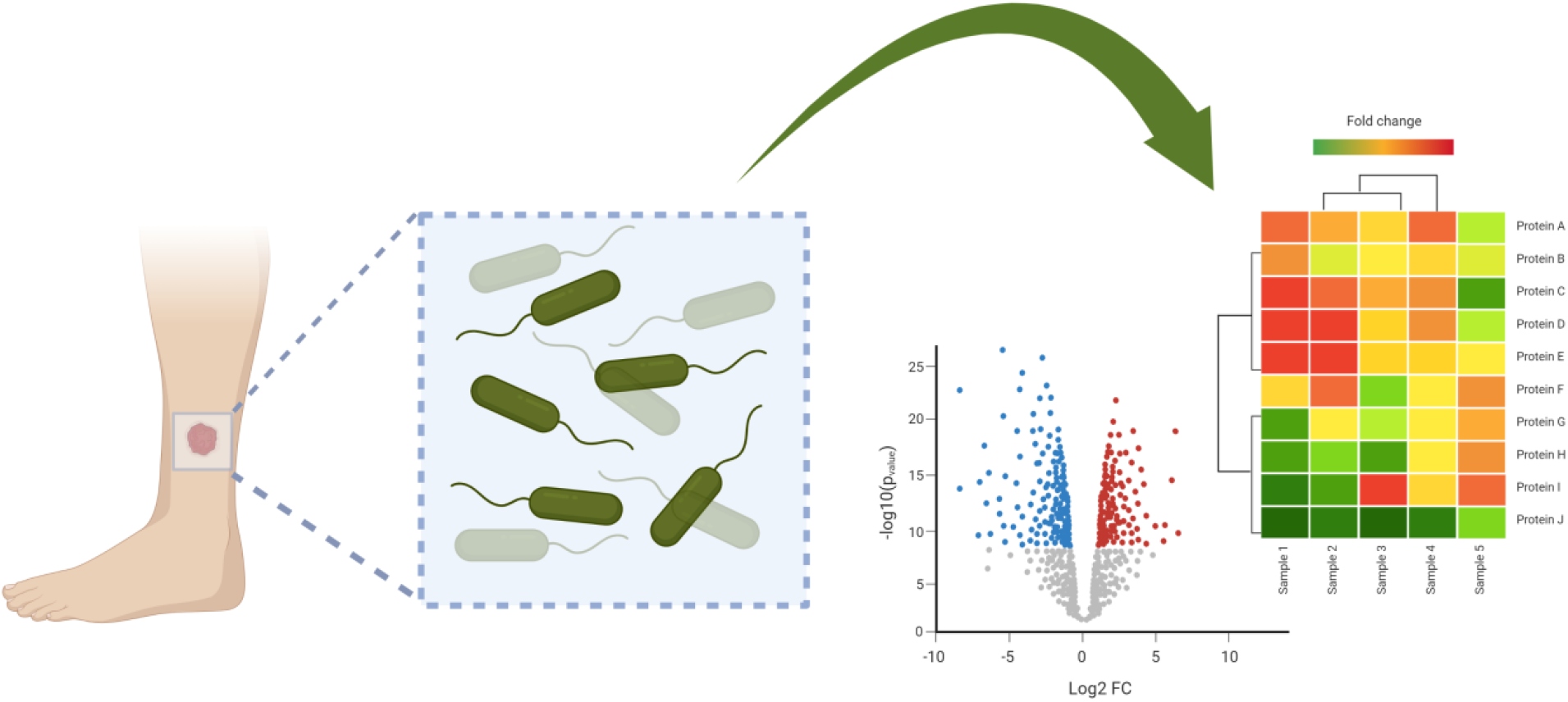
Proteome investigation of mode of action of tobramycin in drug tolerant clinical wound isolates

