## Supplementary tables and figures for "Analysis of antibiotic response in Clinical Wound *Pseudomonas aeruginosa* isolates: Unveiling Proteome Dynamics of tobramycin tolerant phenotype"

**Materials and Methods**

**Minimum inhibition concentration**

The minimum inhibitory concentration (MICs) for the clinical wound isolates and PAO1 towards tobramycin was determined following standard MIC protocols described before (Mojsoska et al., 2015). Briefly, the MIC values were determined by incubating the culture for 24 hours at 37°C, (n=3) with various concentrations (tobramycin 32 - 0.05 µg/ml; the inhibitory effect was observed based on the lack of turbidity, Table S1). Following the MIC, when the would isolates and PAO1 were treated with tobramycin concentrations at 10 x MIC (killing assay), colony forming units (CFU assay) were counted to estimate bacterial survival (Figure S1). The experiment was performed using cationic adjusted Mueller-Hinton Borth (MHBII) (Becton and Dickinson) was used to test the susceptibility to tobramycin.

Table S1. Minimum inhibitory concentration of 3 *Pseudomonas aeruginosa* isolates.

| Isolate | PAO1 | 9000a (patient1) | 989a (patient2) |
| --- | --- | --- | --- |
| MIC (Tobramycin) | 0.4 µg/ml | 0.4 µg/ml | 0.4 µg/ml |

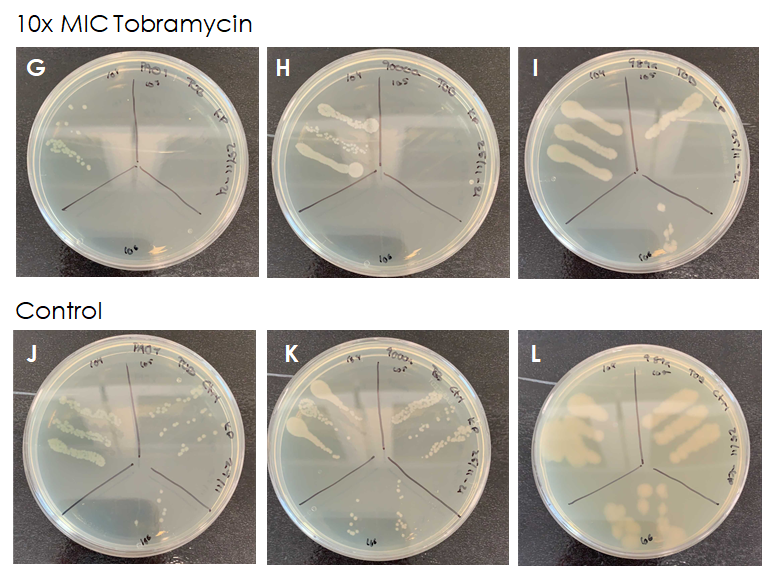

Figure S1. Colony forming units of tobramycin tolerant test. Two *P. aeruginosa* wound isolates 9000a, and 989a and laboratory standard PAO1 (n = 3), were used for the experiment. The overnight culture was diluted 1:100 in 20 ml cationic adjusted Mueller-Hinton Broth (MHBII) and incubated for 24 hours at 37C°. At the stationary phase, the cultures were split into a control group and a treatment group. The culture was treated with 10xMIC tobramycin. After 24 hours of exposure, the culture was plated on LB agar plate for growth assessment. G) PAO1, H) 9000a, I) 989a, J) PAO1 control, K) 9000a, L) 989a, were all cultured in MHBII and treated with tobramycin. All *P. aeruginosa* strains show tobramycin tolerant profile (CFU/ml>10000).

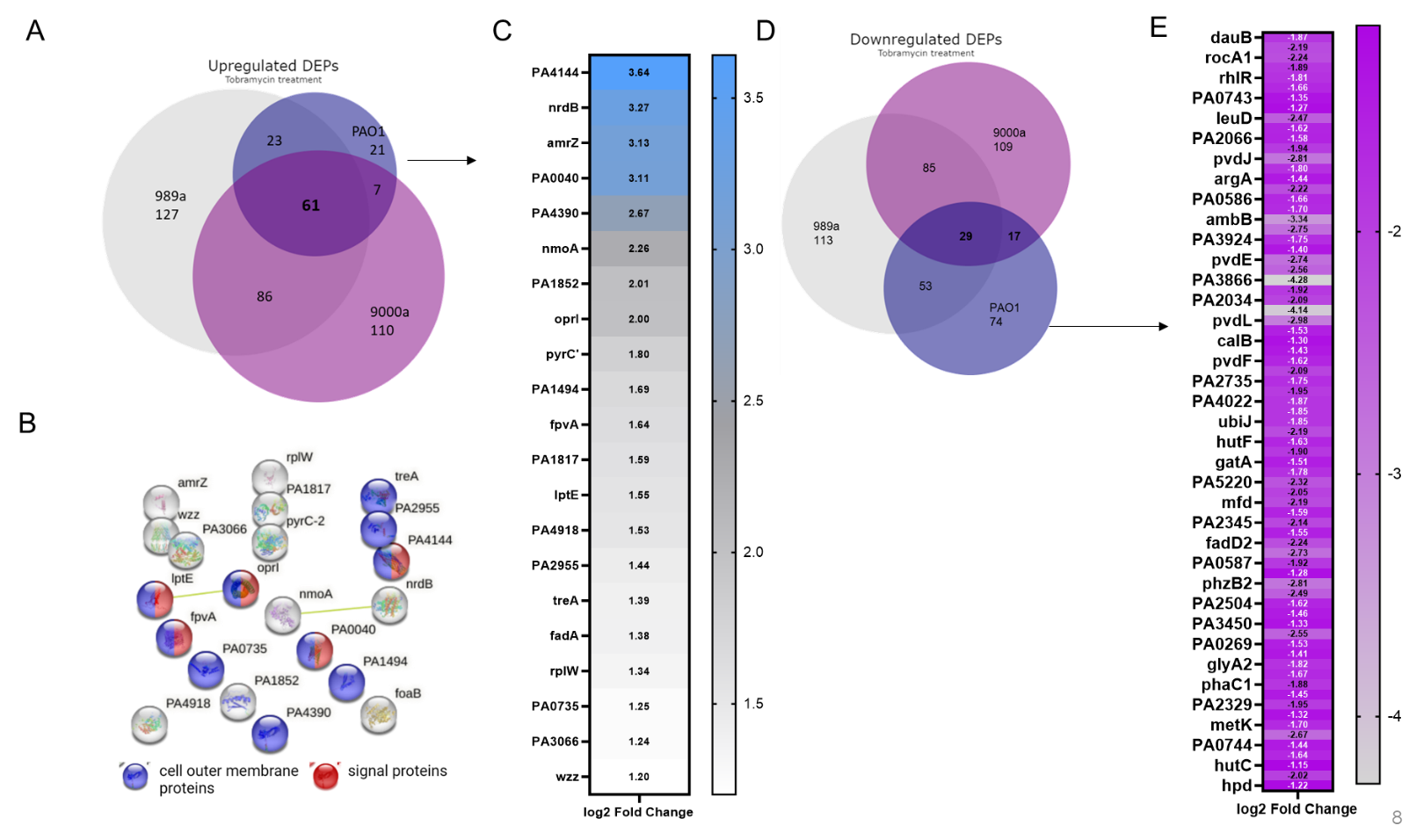

Figure S2. A) Venn diagram of all differentially expressed proteins (DEPs) that are upregulated in the antibiotic treated *P. aeruginosa* isolates, PAO1, 989a and 9000a. B) Visualization and enrichment of the top proteins was done using STRING (version 12. C) Log2-FC of the 21 upregulated DEPs. D) Venn diagram of all differentially expressed proteins (DEPs) that are downregulated in the antibiotic treated *P. aeruginosa* isolates, PAO1, 989a and 9000a. E) Log2-FC of the 74 downregulated DEPs (log2FC>-1.5).

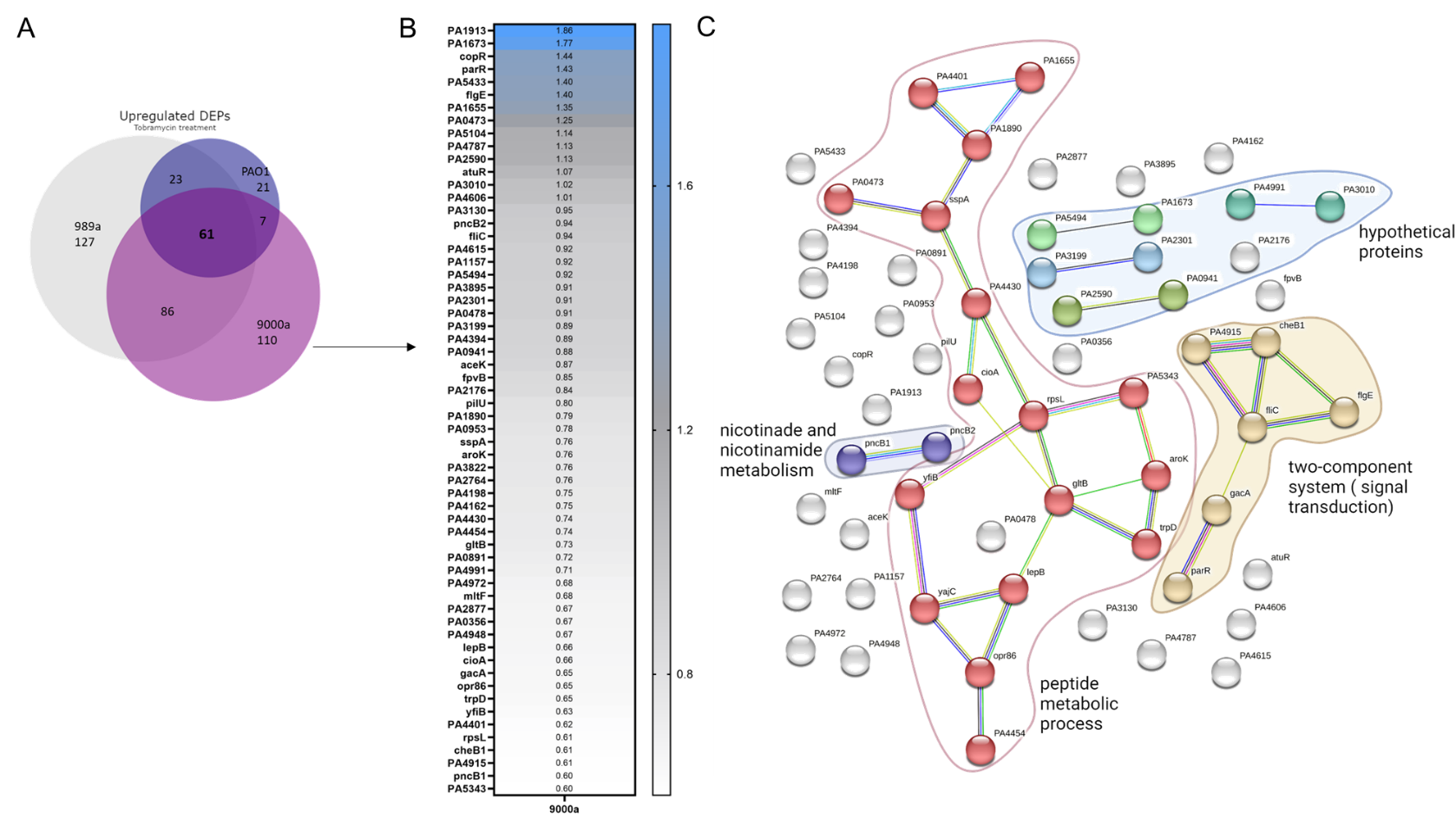

Figure S3. A) Venn diagram of all differentially expressed proteins (DEPs) that are upregulated in the antibiotic treated *P. aeruginosa* isolates, PAO1, 989a and 9000a. B) Log2-FC of DEPs found unique for 9000a with log2-FC >0.5. C) Visualization and enrichment of the top proteins was done using STRING (version 12.0).

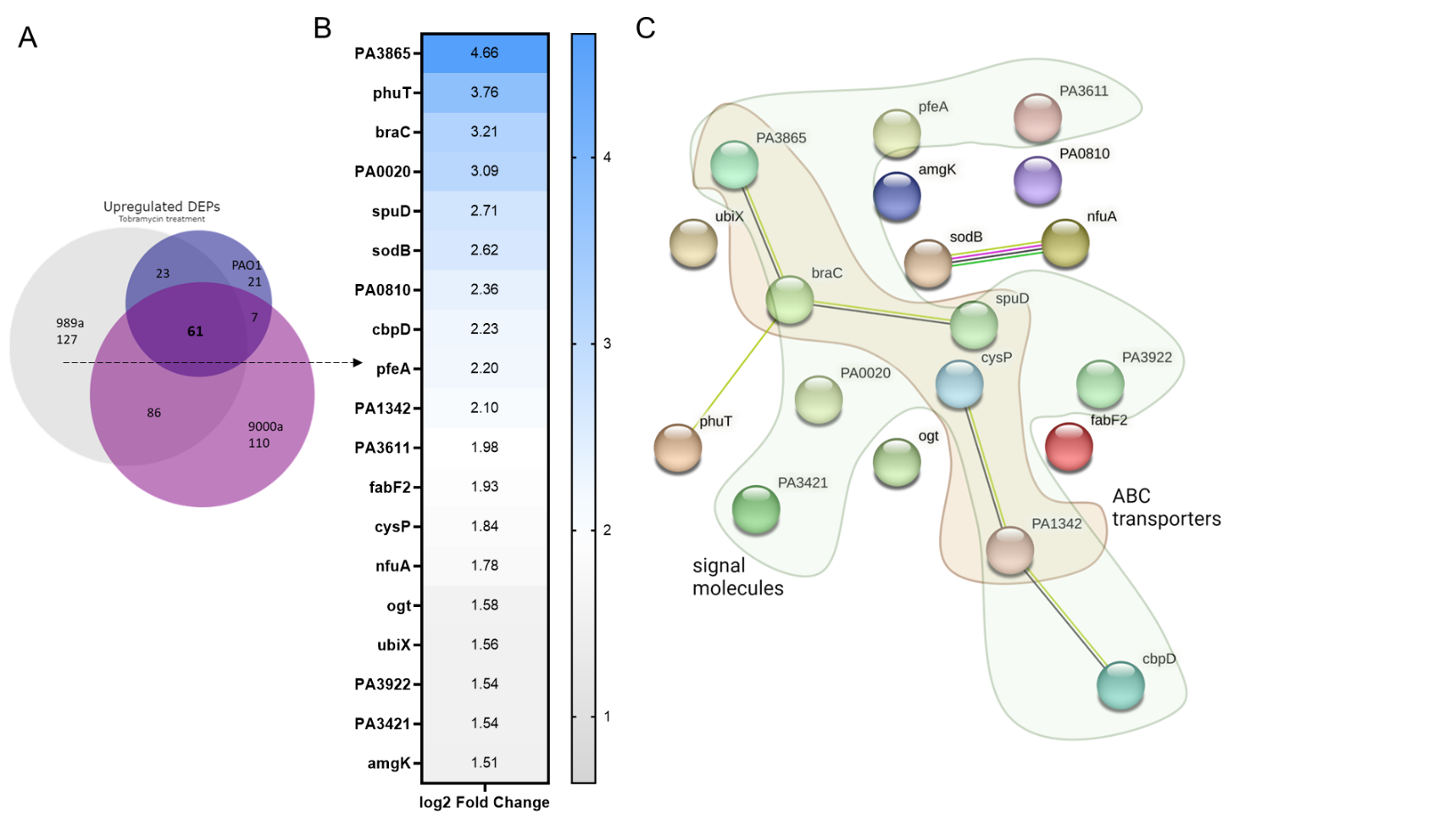

Figure S4. A) Venn diagram of all differentially expressed proteins (DEPs) that are upregulated in the antibiotic-treated *P. aeruginosa* isolates, PAO1, 989a, and 9000a. B) Log2-FC>1.5 of unique DEPs for 989a. C) Visualization and enrichment of the top proteins was done using STRING (version 12.0).
